# PD-1 blockade during T cell priming enhances long-term protection against metastatic tumors by epigenetically tuning T cell exhaustion

**DOI:** 10.1101/2025.11.25.690601

**Authors:** Teresa Dinter, Sebastian Mackowiak, Zachary J. Rogers, Vidit Bhandarkar, Molly Y. Carney, Yiming J. Zhang, Duncan Morgan, Lorelai Pop, Fiona Chatterjee, Emi Lutz, Yajit Jain, Adityanarayanan Radhakrishnan, Eric Lander, K. Dane Wittrup, J. Christopher Love, Alexander Meissner, Stefani Spranger

## Abstract

In cancer, CD8^+^ T cell responses are dominated by exhausted T cells, which can be reinvigorated using immune checkpoint blockade therapy and can control large tumors. However, it remains unclear which T cell fate best supports long-term immunity following tumor regression or clearance and a period of minimal antigen load. This question is particularly relevant following surgical tumor resection, when tuning the immune system could prevent recurrence. To determine which T cell fate provides durable protection following surgery and metastatic rechallenge, we modulated T cell priming using anti-PD-1, IFN-β or agonistic anti-CD40 and assessed effects on CD8^+^ T cell differentiation and overall survival. IFN-β and anti-CD40 promoted effector and memory-like T cell states, respectively, whereas anti-PD-1 did not markedly alter T cell differentiation, yet conferred the greatest survival benefit against metastatic tumors. Notably, anti-PD-1 induced epigenetic remodeling, which was detectable upon metastatic recall, consistent with the maintenance of a circulatory intermediate-exhausted T cell state. Thus, while effector and memory precursor-like T cells could be generated with IFN-β and agonistic anti-CD40, only the intermediate-exhausted T cell state driven by anti-PD-1 supported durable anti-tumor immunity.

**Summary:** This study shows that PD-1 blockade during T cell priming promotes a circulatory intermediate-exhausted CD8⁺ T cell state that uniquely supports durable anti-tumor immunity after surgical resection and metastatic challenge, outperforming effector or memory-like T cell responses generated by IFN-β or CD40 agonist treatment, respectively.

## INTRODUCTION

Most anti-tumor immune responses are largely comprised of exhausted CD8^+^ T cell states. Broadly, exhausted T cells express high levels of inhibitory receptors and exhibit diminished effector function. Despite this, they retain partial cytotoxic capacity and sustain long-lived responses. Therapeutic efforts have therefore focused on reinvigorating exhausted T cells by blocking inhibitory receptors, an approach termed immune checkpoint blockade (ICB) therapy. Specifically, anti-PD-1/PD-L1 blocking antibodies promote the differentiation of progenitor exhausted T cells (T_pex_) into cells with heightened cytotoxic ability capable of mediating tumor clearance (*1–5*).

T cell exhaustion represents one branch of CD8^+^ T cell differentiation, epigenetically distinct from effector and memory lineages found during and following acute infection (*6, 7*). Effector and memory T cell states display robust cytotoxicity and durable persistence, respectively, and have evolved to clear antigen and prevent recurrence. While it has been suggested that some qualities of these cells may enhance anti-tumor immunity, it remains unclear whether steering T cell differentiation toward these states would yield more durable protection compared to ICB-induced exhausted states.

Recent advances in the treatment landscape, such as neoadjuvant ICB therapy or therapeutic vaccination in the adjuvant setting, continue to rely on ICB agents that were originally developed and studied under conditions of high tumor burden and persistent antigen. However, these newer treatment modalities function in settings where the primary tumor has been resected and antigen levels are low, with de novo priming potentially being induced by surgery or therapeutic vaccination. It should therefore be carefully revisited which T cell state would be best suited to establish long-term immunity against metastatic relapse.

New data propose that CD8⁺ T cell fate is established during T cell priming (*8–10*). Thus we tested whether clinically-relevant immunotherapies administered during T cell priming could reshape T cell state and improve outcomes following surgery and metastatic rechallenge in a murine melanoma model. We evaluated three interventions: interferon-beta (IFN-β), an agonistic anti-CD40 antibody, and an antagonistic anti-PD-1 antibody. IFN-β is a cytokine that is highly expressed during acute infections and has been associated with generating cytotoxic effector T cells (*11*). Agonistic anti-CD40 antibody enhances the costimulatory function of dendritic cells and promotes memory T cell responses in some tumor settings (*12, 13*). Anti-PD-1, the current clinical standard, targets T_pex_ and intermediate-exhausted T cells (T_int_) to promote their proliferation and differentiation (*1, 14–17*).

By modulating these pathways during T cell priming, we observed distinct redirection of CD8^+^ T cell differentiation in a murine melanoma model. Strikingly, only anti-PD-1 therapy synergized with surgical tumor resection to enhance protection against metastatic challenge. In contrast, anti-CD40 and IFN-β induced memory and effector states, respectively, and reduced survival of mice compared to surgery alone. Epigenetic profiling of T cells during the recall response indicated that transient blockade of the PD-1 pathway during T cell priming promoted an intermediate-exhausted phenotype. Our findings suggest that the T cell exhaustion differentiation trajectory is indeed optimal for long-term tumor control and that retention of T cells in an intermediate state poises tumor reactive T cells for optimal protection.

## RESULTS

### Altering priming signals changes the phenotype of tumor-reactive CD8^+^ T cells

To generate a preclinical model of anti-PD1-refractory melanoma, cancer cells were isolated from an autochthonous melanoma driven by *BRaf^V600E^*and *Pten^−/-^* (*18*) and transduced with SV40-LargeT antigen, luciferase, and dTomato. This cell line, named Mel1, grows progressively when injected subcutaneously (Fig. 1A). To study tumor-specific CD8^+^ T cell responses, Mel1 cells were engineered to express the model antigen SIYRYGGL (SIY, Mel1.SIY). Mel1.SIY tumors grow similarly to Mel1, though with a modest growth delay (Fig. 1A). In Rag2^−/-^ mice, Mel1.SIY tumors grow significantly larger than in wild-type animals (Fig. 1B), indicating an initial T cell response that ultimately fails to control tumor growth.

**Fig. 1.**
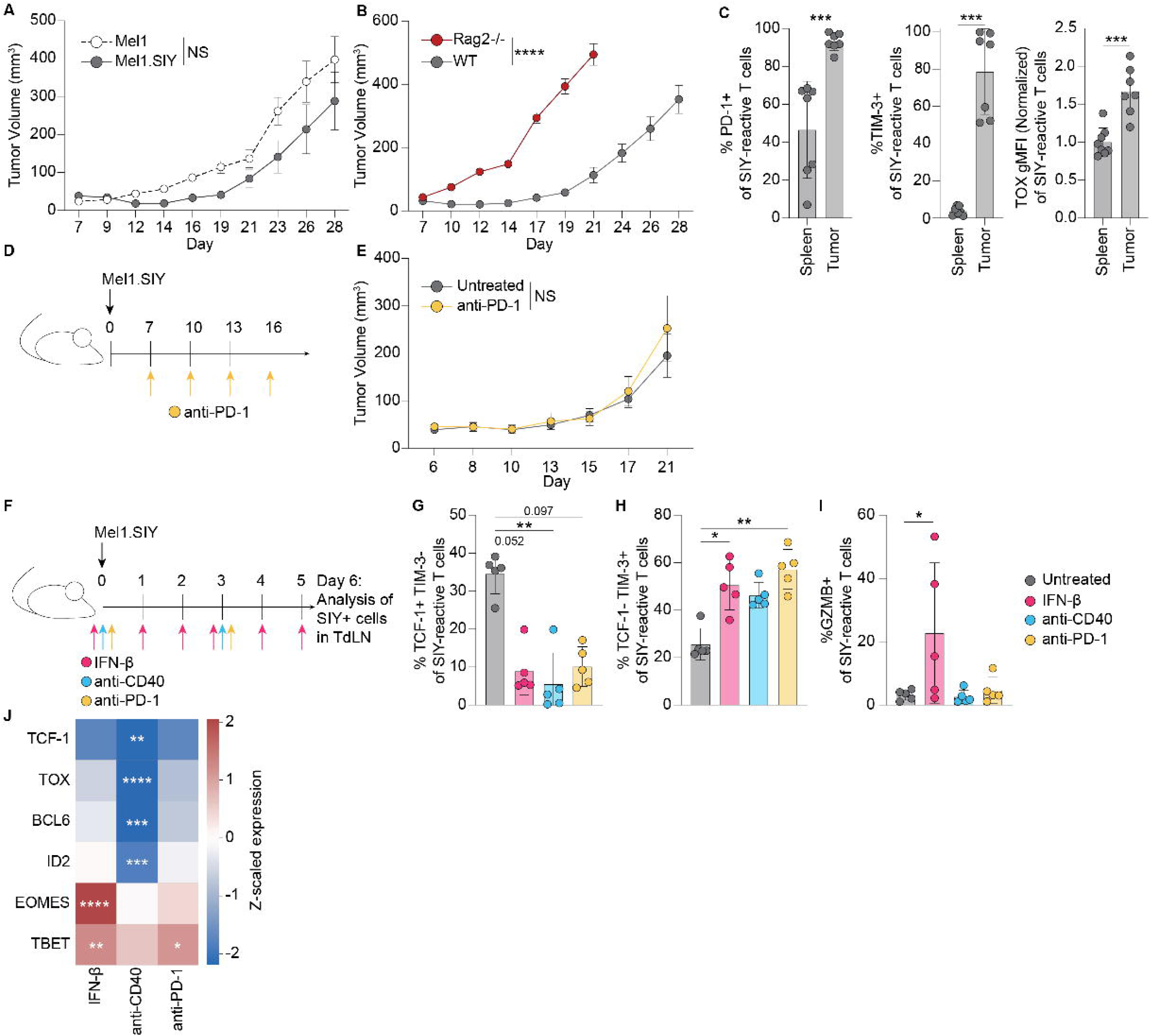
Changing signals during priming against Mel1.SIY tumors alters T cell phenotypes. (**A**) Subcutaneous tumor growth of Mel1 (n=9) and Mel1.SIY (n=5) (**B**) Subcutaneous tumor growth of Mel1.SIY in WT (n=9) and Rag2^−/-^ (n=5) mice. (**C**) Frequency of PD-1^+^ and TIM-3^+^ SIY-reactive T cells in the spleen and tumor. TOX gMFI of SIY-reactive T cells in tumor normalized to average TOX gMFI of SIY-reactive T cells in the spleen. (**D**) Anti-PD-1 treatment schedule for mice bearing Mel1.SIY tumors. (**E**) Subcutaneous Mel1.SIY tumor growth with/without anti-PD-1 treatment (n=8 each). (**F**) Treatment scheme of mice bearing Mel1.SIY tumors. (**G-I**) Frequency of SIY-reactive T cells that are TCF-1^+^TIM-3^−^ (G), TCF-1^−^TIM-3^+^ (H), and GzmB^+^ (I) in the TdLN 6 days after tumor inoculation with treatment. (**J**) Heatmap showing Z-scaled expression relative to untreated mice of SIY-reactive T cells in the TdLN. TCF-1, ID2, EOMES, and TBET are representative of frequency of the indicated gene. TOX and BCL6 are representative of gMFI of the indicated gene. Mean ± sem is shown. p-values calculated using 2-way ANOVA for (A,B,E), Mann Whitney test for (C), Kruskal-Wallis test for (G) and (H), one-way ANOVA for (I). p-value determined for (J) by Kruskal-Wallis test for TCF-1 and TBET and by one-way ANOVA for all others.

To determine whether terminal T cell exhaustion contributed to this failure, we evaluated inhibitory receptor and transcription factors (TF) expression on SIY-reactive T cells. At day 14 post tumor inoculation, tumor-infiltrating T cells expressed high levels of programmed cell death (PD-1), T immunoglobulin and mucin-domain containing-3 (TIM-3), and thymocyte selection-associated high mobility group box (TOX) (Fig. 1C), a TF necessary for establishing and maintaining exhausted T cells (*19, 20*). Since melanoma with exhausted T cells can be responsive to ICB therapy, we evaluated anti-PD-1 therapy in the Mel1.SIY tumor model (Fig. 1D). Interestingly, mice bearing Mel1.SIY tumors failed to respond to anti-PD-1 therapy (Fig. 1E). In sum, we generated a transplantable melanoma model in which T cells become exhausted but are resistant to anti-PD-1 therapy.

Because exhaustion programs can be imprinted early during T cell priming (*9, 10*), we characterized the kinetics of this phase, analyzing the number, frequency, and phenotype of SIY-reactive T cells in the tumor-draining lymph node (TdLN) post tumor implantation (Fig. S1A). SIY-reactive T cells were first apparent at day 4, plateauing between days 6 to 8, with all SIY-reactive T cells expressing CD44 by day 6 (Fig. S1B-E). Additionally, PD-1 expression was highest on days 5 and 6 (Fig. S1F), suggesting maximal T cell receptor (TCR) stimulation in the TdLN during this window. Thus, day 6 was selected for subsequent analyses as the peak of T cell priming.

To test whether modulating priming signals could alter T cell fate, we chose three distinct interventions; IFN-β promotes robust effector T cell responses (*11*); agonistic anti-CD40 is associated with improved memory T cell responses in some contexts (*13, 21, 22*); and anti-PD-1 improves anti-tumor responses by expanding and differentiating T_pex_ (*1, 16, 17*). To understand how treatment during T cell priming impacts T cell fate and the resulting anti-tumor immune response, we treated mice with either IFN-β, anti-CD40, or anti-PD-1 at the time of tumor inoculation (Fig. 1F). Antibody therapies were dosed again at day 3, and IFN-β was dosed daily due to its shorter half-life (Fig. 1F) (*23*). Anti-CD40 treatment increased the number of SIY-reactive T cells (Fig. S1G). All therapies promoted differentiation of SIY-reactive T cells from TCF-1^+^ TIM-3^−^ T_pex_ to TCF-1^−^ TIM-3^+^ T cells (Fig. 1G-H, Fig S1H-I). IFN-β treatment significantly increased granzyme B (GzmB) expression (Fig. 1I), consistent with enhanced effector T cell generation.

Because T cell differentiation is controlled by several TFs (*20, 24–34*), we evaluated their expression following treatment (Fig. S1J-O). All interventions decreased TOX expression, which was most substantial with anti-CD40 treatment (Fig. 1J, Fig. S1K). Anti-CD40 treatment also decreased expression of BCL6 and ID2 (Fig.1J, Fig. S1L-M), which promote memory and stem-like features and regulate the balance between effector and exhausted phenotypes, respectively (*27–30*). IFN-β treatment strongly induced expression of EOMES and, to a lesser extent, TBET (Fig. 1J, Fig. S1N-O). In contrast, anti-PD-1 selectively upregulated TBET (Fig. 1J, Fig. S1O). A prior study demonstrated that the relative balance of these TFs is critical, with higher EOMES expression associated with exhausted T cell and elevated TBET levels were linked with memory T cells (*34*). These data establish that altering signals during T cell priming results in phenotypic changes and altered TF expression, suggesting the potential of these therapies to alter CD8^+^ T cell fate.

### Anti-PD-1 therapy maintains T cell phenotypes similar to unperturbed priming, while IFN-β and anti-CD40 treatment create effector and memory-like T cells, respectively

To gain an unbiased understanding of the transcriptional changes induced by altering priming signals, we performed paired single-cell RNA (scRNA-seq) and TCR sequencing of SIY-reactive T cells from TdLNs at days 4 and 6 post tumor inoculation to capture T cells during early and peak priming using Seq-Well (*35*). After removing low-quality cells, we used Unifold Manifold Approximation and Projection (UMAP) for dimensionality reduction and identified 9 clusters (Fig. 2A). Cluster identities were based on top differentially expressed genes and included early activated cells, two proliferative clusters, two effector clusters, one memory precursor cluster, two progenitor exhausted clusters, and an interferon stimulated gene (ISG) cluster (Fig. 2A-C). Notably, T cells in the memory precursor cluster expressed genes associated with resident memory T cells (Trm) (Fig. 2B-C). Progenitor exhausted 2 cluster T cells expressed canonical exhaustion markers, including *Lag3*, *Tcf7*, *Slamf6*, and *Pdcd1* (Fig. 2B-C), while T cells in the progenitor exhausted 1 cluster expressed a noncanonical set of exhaustion-associated genes such as *Myb*, *Hells*, and *Mt1* (Fig. 2B-C).

**Fig. 2.**
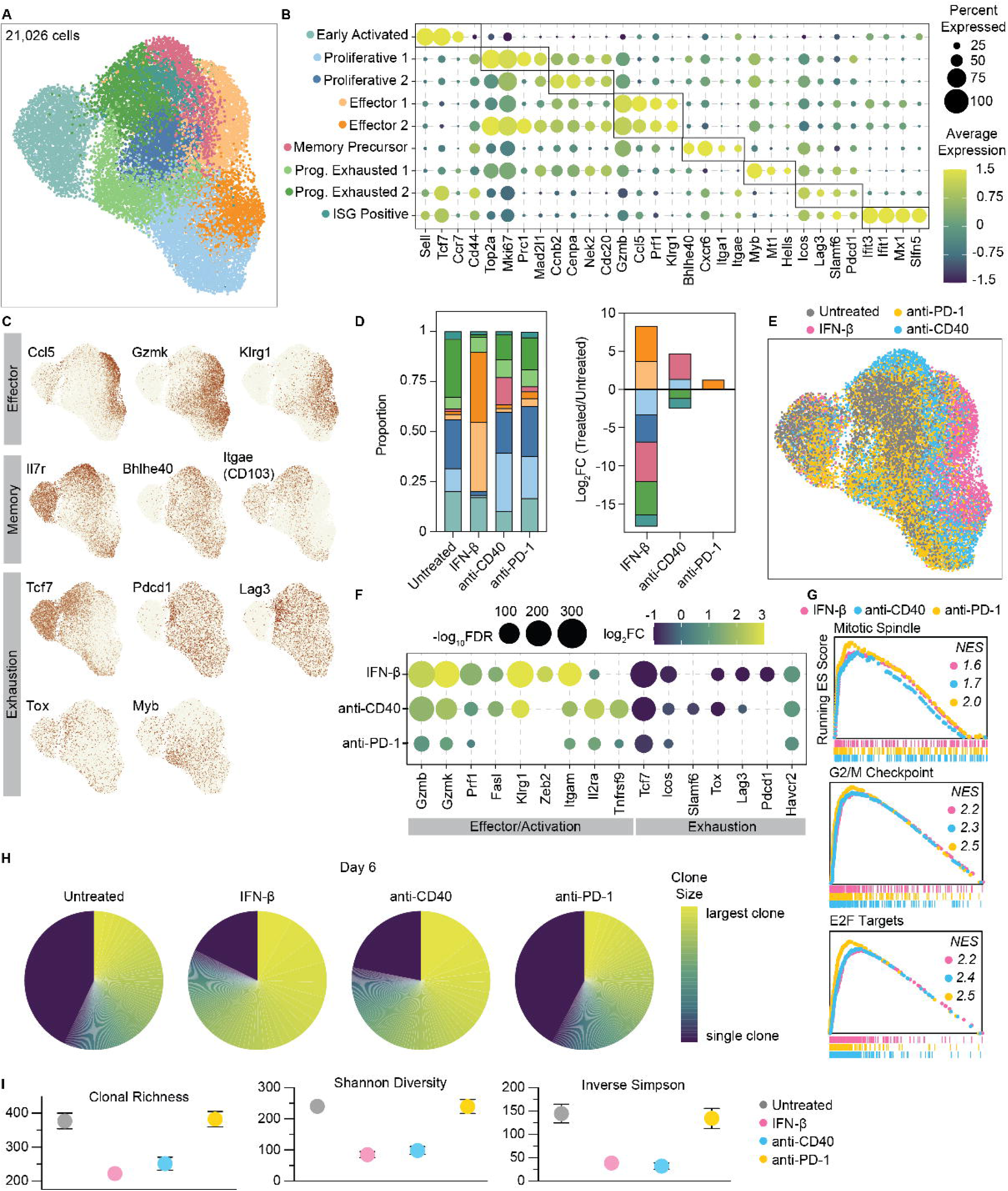
Anti-PD-1 therapy increases clonal expansion while maintaining T cell phenotypes similar to native priming, while IFN-β and anti-CD40 treatment create effector and memory-like cells, respectively. (**A**) scRNA-seq UMAP of 21,026 SIY-reactive T cells from the TdLN colored by T cell phenotype clusters. (**B**) Dot plot of selected differentially expressed genes used to identify clusters. (**C**) Feature plots showing expression for selected genes on projected UMAP. (**D**) Proportion of each cluster’s contribution to the indicated treatment group. (**E**) UMAP of SIY-reactive T cells from TdLN colored by treatment group. (**F**) Differential gene expression analysis of effector or exhaustion related genes for each treatment group relative to untreated controls. (**G**) GSEA analysis of indicated Hallmark cell cycle gene sets. (**H**) Top 100 TCR clones at day 6, wherein each slice represents one clone ranked by the largest clone to smallest, shown for each treatment group. (**I**) Diversity indices of TCR clonotypes for each treatment group, including Inverse Simpson, clonal richness, and Shannon diversity for treatments at day 6.

We next asked how each intervention skewed T cell phenotype. In untreated mice, T cells primarily occupied early activated, proliferating, and progenitor exhausted 2 clusters (Fig. 2D-E), consistent with a canonical exhaustion trajectory. All therapies decreased the proportion of progenitor exhausted 2 cells (Fig. 2D). Strikingly, the majority of IFN-β-treated T cells were in effector clusters, with very few T cells in proliferating clusters (Fig. 2D-E). Anti-CD40 treated T cells uniquely occupied the memory precursor cluster in addition to proliferating and progenitor exhausted clusters (Fig. 2D-E). In contrast, anti-PD-1-treatment maintained a similar distribution seen in untreated controls (Fig. 2D-E).

We next compared effector and exhaustion-associated gene expression relative to the untreated control. IFN-β induced robust upregulation of effector markers, while anti-PD-1 showed minimal transcriptional changes (Fig. 2F). Notably, *Tcf7* expression was decreased among all treatments relative to untreated controls, consistent with flow cytometric analyses (Fig. 1J, Fig. 2F, Fig. S1J). Although anti-CD40 treatment led to the largest increase in SIY-reactive T cell numbers (Fig. S1G), GSEA analysis revealed enrichment in cell cycle genes in all treatment groups, with the greatest enrichment among anti-PD-1-treated T cells (Fig. 2G). Together, these data suggest that IFN-β signaling during T cell priming promotes an effector T cell program, CD40 agonism generates a memory precursor population with features of Trm, and anti-PD-1 treatment maintains T cells along the exhaustion trajectory while enhancing proliferation.

To explore the temporal dynamics of T cell differentiation, we analyzed the day 4 and day 6 datasets separately (Fig. S2A). As expected, most of the day 4 T cells occupied the early activated cluster (Fig. S2A). However, both anti-CD40 and anti-PD-1 increased the proportion of T cells within the proliferative and progenitor exhausted clusters at this early timepoint (Fig. S2B), indicating accelerated T cell differentiation. Supporting this, global transcriptome analyses showed that at day 4, anti-CD40 and anti-PD-1-treated T cells were transcriptionally most similar to each other (Fig. S2C). By day 6 however, the transcriptome of anti-PD-1 treated T cells most closely resembled untreated controls (Fig. S2C). Together, these findings show that while both anti-CD40 and anti-PD-1 therapy accelerates T cell differentiation, only anti-PD-1 maintains T cells within the exhaustion trajectory.

Because ICB treatment is often associated with changes in T cell clonotypes (*36–39*), we examined whether therapy during T cell priming affected the T cell repertoire. In all conditions, T cell clones expanded from day 4 to day 6 (Fig. 2H, Fig. S2D). At day 4, all therapies increased clonal expansion compared to unperturbed priming (Fig. S2D), while at day 6 IFN-β and anti-CD40 groups continued to show enhanced clonal expansion with IFN-β having the most pronounced impact (Fig. 2H). Correspondingly, TCR repertoire diversity metrics decreased in IFN-β and anti-CD40-treated groups (Fig. 2I, Fig. S2E). We next examined clonal overlap between treatments and found the highest overlap between anti-CD40 and anti-PD-1 treated T cells at day 4, while untreated T cells and T cells treated with anti-PD-1 showed highest similarity at day 6 (Fig. S2F-G), mirroring RNA transcriptomic relationships. Among the top 100 most abundant TCR clones, some clonotypes were shared between all groups (Fig. S2H-I). These data indicate that IFN-β and anti-CD40 treatment during priming affects TCR repertoire diversity and expansion, while anti-PD-1 treatment does not significantly alter clonal expansion from unperturbed priming.

Together, these data support that treatment during priming imparts distinct transcriptional and clonal profiles on CD8⁺ T cells. IFN-β drives a strong effector-like program and robust expansion of a limited set of TCR clones. In contrast, anti-CD40 promotes a unique memory precursor state accompanied by similar clonal expansion. Anti-PD-1 therapy reduces expression of the stem-like marker Tcf7, while maintaining a transcriptome and TCR repertoire most like unperturbed priming.

### Anti-PD1 therapy during priming provides therapeutic benefit following surgery and metastatic challenge

Next, we aimed to directly test whether the treatment-induced shifts in T cell fate translate into improved protection from metastatic rechallenge. To do this, tumor-bearing mice received one of the three therapies during priming, after which Mel1.SIY flank tumors were surgically resected, and the mice rechallenged intravenously (IV) with the same cell line 5-8 weeks later without further treatment (Fig. 3A). Tumors following IV-inoculation primarily localized to the lungs and subcutaneous tissue, particularly at the neck and axilla (Fig. S3A), mirroring the metastatic pattern observed in patients with melanoma. Interestingly, mice with prior flank tumors developed fewer subcutaneous lesions upon rechallenge (Fig. S3A), suggesting tissue-specific memory responses. While this model does not replicate the clinical treatment scenario, it enables controlled manipulation of T cell priming to address whether altering T cell fate affects the anti-tumor immune response following surgery and metastatic recurrence.

**Fig. 3.**
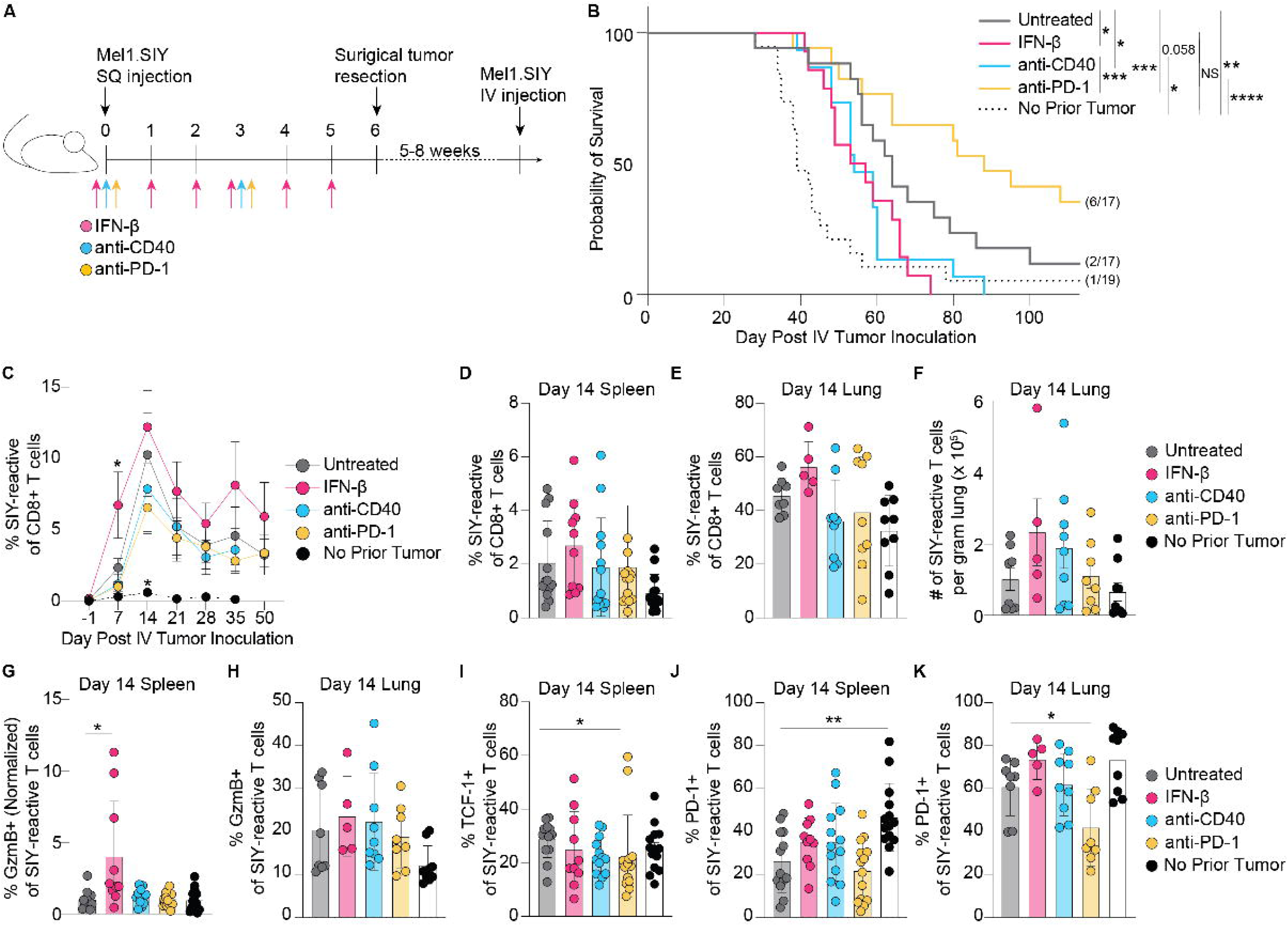
Anti-PD-1 during priming extends survival following metastatic rechallenge. (**A**) Experimental approach in which therapy is administered during priming, tumors are surgically removed, and 5-8 weeks later mice are challenged with IV-inoculation of Mel1.SIY tumors. (**B**) Kaplan-Meier survival curve of mice after IV-rechallenge with Mel1.SIY for untreated (n= 17), IFN-β (n= 14), anti-CD40 (n= 15), anti-PD-1 (n= 17), no prior flank tumor(n= 19) (**C**) Frequency of SIY-reactive T cells of total CD8^+^ T cells in the blood after IV-rechallenge (n= 4), IFN-β (n= 4), anti-CD40 (n= 3), anti-PD-1 (n= 5), no prior flank tumor (n= 5) (**D**) Frequency of SIY-reactive T cells in the spleen 14 days after IV Mel1.SIY inoculation. Quantification of the (**E**) frequency and (**F**) number of SIY-reactive T cells in the lung. Quantification of GzmB^+^ SIY-reactive T cells in the spleen (**G**) and lungs (**H**). GzmB expression in the spleen was normalized to untreated controls. (**I**) Frequency of TCF1^+^ of SIY-reactive T cells in the spleen. Frequency of PD-1^+^ of SIY-reactive T cells in the spleens (**J**) and lungs (**K**) of mice 14 days after IV Mel1.SIY inoculation. Mean ± sem is shown. P-value calculated by log-rank (Mantle-Cox) test for (B), by one-way ANOVA for each timepoint in (C) and for (E,J) and by Kruskal-Wallis test for (E-I,K).

Control mice without prior flank tumors had the shortest median survival of 39 days while mice undergoing surgery without additional treatment (untreated) survived a median of 64 days (Fig. 3B, Fig. S3B), emphasizing the importance of a pre-existing T cell response. Surprisingly, despite inducing memory-precursor-like transcriptional profiles at the time of tumor resection (Fig. 2C), anti-CD40 treatment during priming significantly reduced survival relative to untreated controls (Fig. 3B). Similarly, IFN-β during priming failed to improve the overall survival (Fig. 3B), despite generating effector-like T cells (Fig. 2C). In contrast, anti-PD-1 significantly outperformed all other treatment groups, extending median survival to 88 days (Fig. 3B, Fig. S3B). Compared to mice that had no prior tumor, anti-PD-1 during priming provided greater therapeutic benefit (p-value <0.0001) than surgery alone (p value = 0.0029), with 35% (6/17) of mice achieving long-term survival versus 12% (2/17) in untreated controls. Notably the survival benefit conferred by anti-PD-1 in this setting parallels clinical results in trials of neoadjuvant ICB in melanoma (*40, 41*).

To understand the basis of the survival benefit observed in groups with exhausted T cells (untreated and anti-PD-1), we assessed the magnitude of the SIY-reactive T cell response in the blood during metastatic rechallenge (Fig. S3C). The frequency of SIY-reactive T cells was much greater in mice with prior tumors, consistent with a recalled immune response (Fig. 3C). Interestingly, IFN-β treatment yielded the highest immediate frequency of SIY-reactive T cells in the blood (Fig. 3C), suggesting that rapid expansion of systemic T cell responses alone is not protective.

We next profiled SIY-reactive T cells at day 14 after metastatic challenge, corresponding to the peak of the circulating response, examining T cells in the white pulp of the spleen and in lung parenchyma to assess systemic and tumor-site T cell responses (Fig. S3D-E). Across both tissues, the frequency and number of SIY-reactive T cells were comparable across all treatment groups (Fig. 3D-F). IFN-β treatment increased GzmB expression in splenic SIY-reactive T cells, but this was not observed in the lungs (Fig. 3G-H). In the spleen, TCF-1 expression was modestly reduced in SIY-reactive T cells from mice treated with anti-PD-1 (Fig. 3I), suggesting fewer T_pex_ at this timepoint. PD-1 expression on splenic T cells was unchanged with therapy (Fig. 3J). However, T cells from the lungs of anti-PD-1-treated mice retained lower PD-1 expression (Fig. 3K), potentially implying reduced sensitivity to inhibitory cues or lower antigen burden within the lung microenvironment. Together these data suggest that the quality of persisting T cells during metastatic recall is shaped by priming signals. Remarkably, only two doses of anti-PD-1 during priming conferred durable benefits over 100 days later, extending survival and limiting PD-1 upregulation on tumor-reactive T cells in the tumor microenvironment.

### Blockade of PD-1 during T cell priming improves both effector function and memory potential

To assess the relative contributions of cytotoxic ability versus memory potential to the efficacy of anti-PD-1 administered during priming, we used an adoptive transfer approach to isolate CD8^+^ T cell-intrinsic effects. Naïve congenically-marked TCR-transgenic 2C T cells, specific for SIY presented on H2-K^b^, were transferred into tumor-bearing mice and treated throughout their priming window (Fig. S4A). Four days later, 2C T cells from TdLNs were analyzed and confirmed to reflect the endogenous response (Fig. S4A-B). All treatments promoted differentiation, reducing the frequency of TCF-1^+^TIM-3^−^ T_pex_ and increasing frequency of TCF-1^−^TIM-3^+^ T cells, with IFN-β treatment reaching statistical significance (Fig. S4C-E). IFN-β also increased GzmB expression (Fig. S4F). TF expression in transferred 2C T cells largely mirrored the endogenous response (Fig. 1N, Fig. S4G-K). Importantly, this approach yielded substantially more T cells in TdLNs compared to the endogenous response, enabling subsequent adoptive transfers (Fig. S4L-M).

To assess cytotoxic ability, in vivo-primed 2C T cells exposed to the different therapies were isolated from TdLNs, and 20,000 T cells were then transferred into treatment-naïve, tumor-bearing Rag2^−/-^ mice, which were subsequently monitored for tumor growth (Fig. 4A). All groups receiving 2C T cells showed delayed tumor growth relative to mice without transferred T cells (Fig. 4B). However, only anti-PD-1-treated T cells significantly slowed tumor progression compared to untreated control T cells (Fig. 4B). Surprisingly, despite high GzmB expression (Fig. S4F), IFN-β-treated T cells did not improve tumor control (Fig.4B). To confirm the diminished cytotoxicity of IFN-β-treated T cells, we performed an in vivo killing assay. A 1:1 mixture of SIY-loaded (CellTrace Violet-labeled) and unloaded (CFSE-labeled) CD45.1^+^ splenocytes were transferred into tumor-bearing CD45.2^+^ mice that were treated with therapy during T cell priming (Fig. 4C). 24 hours later, we analyzed the ratio of SIY-loaded to unloaded splenocytes recovered from the spleen. Mice treated with IFN-β or anti-CD40 exhibited significantly reduced killing of SIY-loaded targets compared to untreated controls (Fig. 4D), confirming impaired cytotoxic function under these treatment conditions. In contrast, anti-PD-1-treated and control mice showed efficient killing (Fig. 4D).

**Fig. 4.**
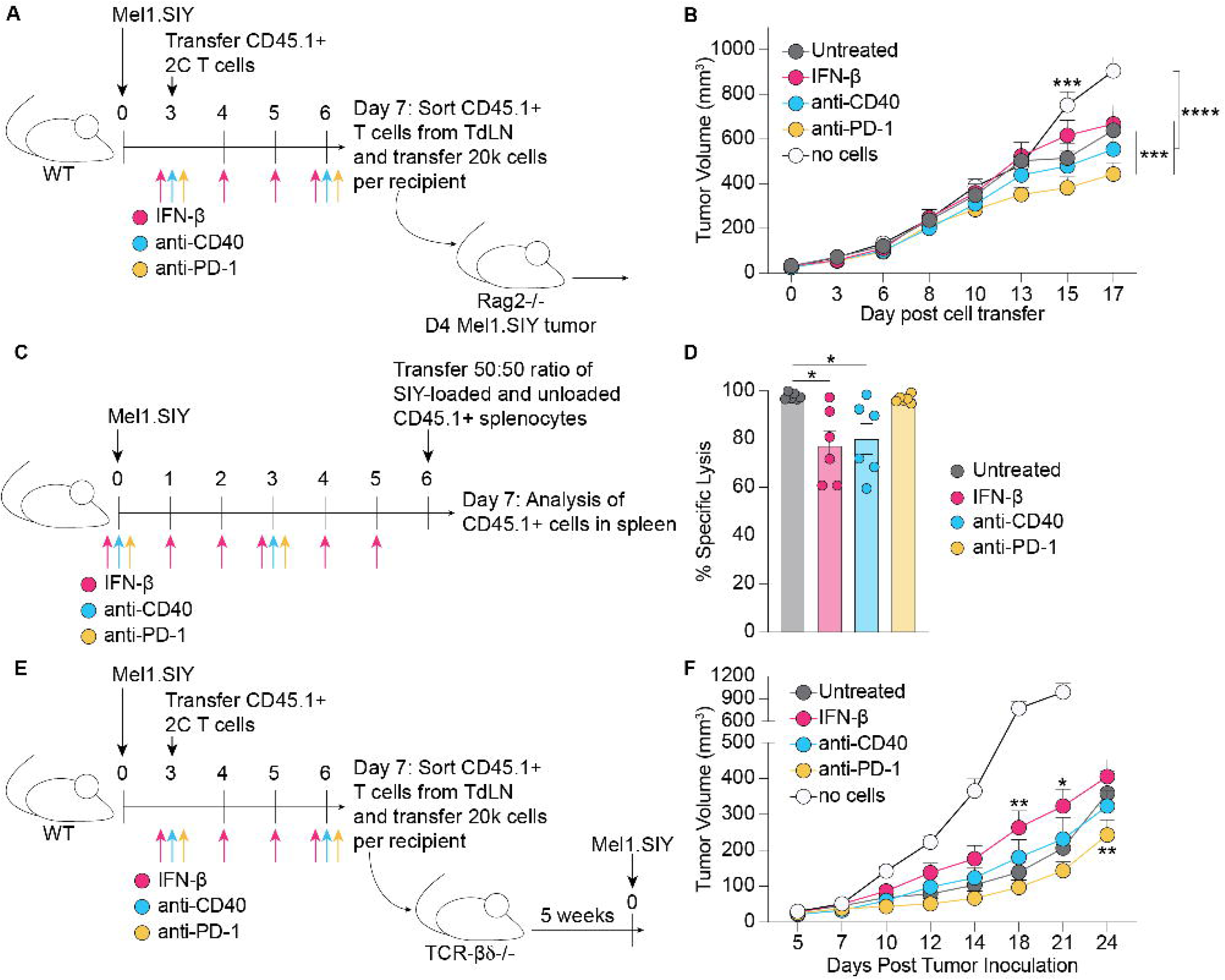
PD-1 blockade during priming improves effector function and memory potential. (**A**) Strategy to access effector function in an adoptive transfer approach. (**B**) Tumor outgrowth of Rag2^−/-^ mice in (A). (**C**) Experimental design for in vivo cytotoxicity assay. (**D**) Quantification of specific lysis of congenically-labeled cells recovered from the spleen. (**E**) Adoptive transfer strategy to access memory potential of T cells treated during priming. (**F**) Outgrowth of tumors in TCRβδ^−/-^ mice from (E). Mean ± sem is shown. p-values calculated using 2-way ANOVA for (B) and (F) and one-way ANOVA for (D).

Next, we evaluated memory potential using a similar adoptive transfer approach. Treated, in vivo-primed 2C T cells from TdLNs were transferred into TCR-βδ^−/-^ mice, rested for 5 weeks to allow for memory formation, and then challenged with subcutaneous Mel1.SIY tumors (Fig. 4E). Expectedly, tumors grew rapidly in mice without transferred 2C T cells (Fig. 4F). IFN-β treatment during T cell priming worsened tumor control, suggesting that skewing T cells toward an effector-like state impairs recall capacity (Fig. 4F). Only mice receiving in vivo-primed 2C T cells treated with anti-PD-1 exhibited delayed tumor growth compared to untreated controls (Fig. 4F). These findings indicate that PD-1 blockade during priming elicits durable cell-intrinsic effects, enhancing long-lasting cytotoxic potential more than 8 weeks after the final dose.

### Anti-PD-1 during T cell priming promotes an intermediate exhausted epigenetic state in the spleen

Given that functional differences persisted more than 5 weeks after treatment, we hypothesized that epigenetic changes might lead to functionally distinct T cell states. To examine how anti-PD-1 during T cell priming influences T cell differentiation, we performed single-cell ATAC sequencing of SIY-reactive T cells across multiple timepoints. T cells were isolated from TdLNs on day 6 post subcutaneous tumor inoculation, representative of the peak of T cell priming, and from the spleen and lung 14 days after metastatic rechallenge (>D59 post-flank tumor) (Fig. 5A). After quality control, cells were projected into UMAP space (Fig. 5B). Clustering was primarily driven by tissue site, with chromatin accessibility from the same tissue showing the highest similarity (Fig. 5B-C).

**Fig. 5.**
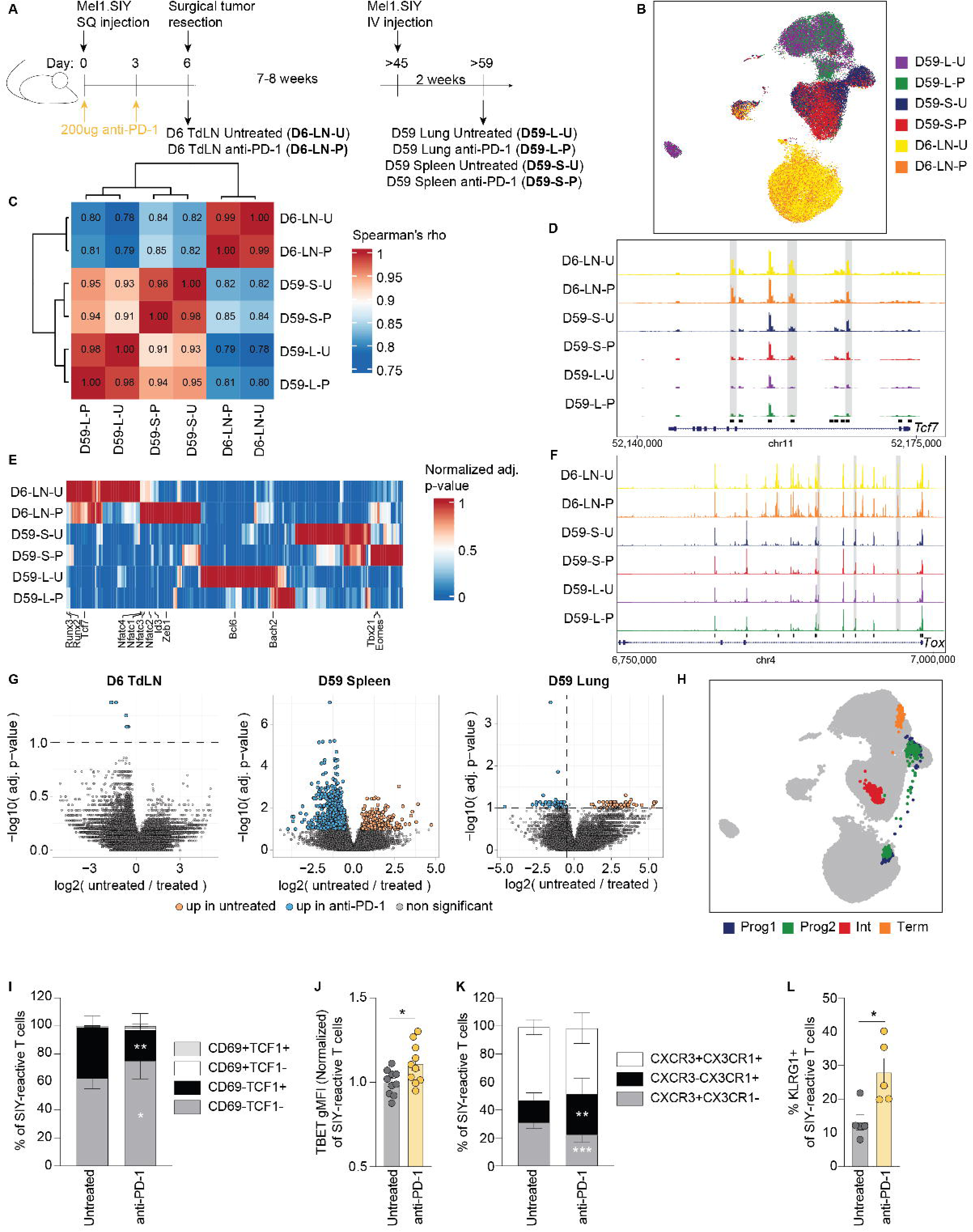
Anti-PD-1 during T cell priming programs durable epigenetic changes consistent with altered T cell exhaustion differentiation. (**A**) Experimental approach to generate samples for single-cell ATAC sequencing. (**B**) UMAP colored by sample identity. (**C**) Spearman analysis of global similarity between ATAC samples. (**D**) ATAC signal tracks of the *Tcf7* locus. (**E**) Analysis of transcription factor motifs in differentially accessible regions. (**F**) ATAC signal tracks of the *Tox* locus. (**G**) Pairwise comparisons of differentially accessible regions for each tissue comparing untreated to anti-PD-1-treated samples. (**H**) UMAP of (B) overlaid with T cell exhaustion subsets from published data (*45*). (**I**) Flow cytometry of CD69 and TCF1 expression of SIY-reactive T cells isolated from spleens 14 days after metastatic challenge. (**J**) TBET expression of SIY-reactive T cells isolated from spleens 14 days after metastatic challenge. (**K**) Flow cytometry of CXCR3 and CX3CR1 expression of SIY-reactive T cells isolated from spleens 14 days after metastatic challenge. (**L**) KLRG1 expression of SIY-reactive T cells isolated from spleens 14 days after metastatic challenge. Mean ± sem is shown. p-values calculated using unpaired T test for (I-L).

Next, we analyzed genes associated with exhausted T cell differentiation. Accessibility of the *Tcf7* locus highest in the TdLN and lowest in lung T cells (Fig. 5D), consistent with progressive differentiation. TF motif analysis of differentially accessible regions (DARs) revealed *Tcf7* motif enrichment in D6 TdLN T cells from untreated mice, whereas anti-PD-1-treated D6 TdLN T cells were enriched for *Nfat* family motifs and *Id3*, a TF linked to stem-like T cells and restraining terminal T cell exhaustion (Fig. 5E) (*7, 42–44*). These results suggest that PD-1 blockade shifts early differentiation away from a canonical TCF-1-driven T_pex_ state toward a distinct *Id3*-supported stem-like program. Following metastatic rechallenge, splenic T cells from anti-PD-1-treated mice showed reduced *Tox* accessibility (Fig. 5F), indicating decreased sensitivity to exhaustion-inducing stimuli; however, accessibility at canonical exhaustion loci (*Pdcd1*, *Lag3*, *Havcr2*) remained unchanged.

Unbiased DAR analysis revealed the most differences between untreated and anti-PD-1-treated T cells in the spleen during metastatic challenge (Fig. 5G). In line with this, the spleen was the only tissue in which samples clustered by treatment (Fig. 5B, Fig. S5A-B). Mapping DARs to nearby genes (<10kb), untreated T cells showed increased accessibility at the stress-response genes (*Ubr1*, *Eif2a*) and select memory-associated loci (*Ahr*, *Usp1*), whereas other memory-linked loci (*Bhlhe40*, *Runx2*, *Myb*, *Ets1*) were less accessible (Table S1). Anti-PD-1-treated T cells had increased accessibility at TCR-signaling related genes (*Nfkb1*, *Itk*, *Arap2*, *Nck2*) and costimulatory receptors (*Tnfrsf9, Cd82*) (Table S1), suggesting enhanced capacity for functional restimulation. Consistent with this notion, *Klf2*, a TF linked to limiting terminal T cell exhaustion, was also more accessible in splenic T cells from anti-PD-1-treated mice (Table S1), although motif enrichment for Klf2 was not observed. Anti-PD1 therapy additionally increased *Ctla4* accessibility post-rechallenge (Table S1), suggesting that anti-PD-1 treatment may prime CD8^+^ T cells to be more responsive to subsequent anti-CTLA4 therapy.

To contextualize these states, we overlaid a published bulk ATAC-seq reference dataset of progenitor (Tex^prog1^, Tex^prog2^), intermediate (Tex^int^), and terminal (Tex^term^) exhausted T cell subsets (*45*), each computationally extrapolated into 250 cells. Tex^prog^ overlapped with TdLN and splenic T cells, Tex^int^ with splenic T cells, and Tex^term^ with lung T cells (Fig. 5H), reflecting lymphoid compartmentalization of stem-like populations. In the spleen, Tex^prog^ subsets clustered with untreated T cells, while Tex^int^ clustered with anti-PD-1-treated T cells (Fig. 5H). Subsetting spleen samples and re-overlaying the reference dataset confirmed this pattern (Fig. S5C), suggesting shared epigenetic features between Tex^int^ and T cells treated with anti-PD-1 during priming. The reference study defined subsets by Slamf6 and CD69 expression (*45*). Because TCF-1 expression closely parallels Slamf6 (*2, 45*), we used CD69 and TCF-1 to validate these findings by flow cytometry. Anti-PD-1 treatment during T cell priming reduced CD69^−^TCF-1^+^ (Tex^prog2^-like) and expanded CD69^−^TCF-1^−^ (Tex^int^-like) populations (Fig. 5I), confirming our ATAC sequencing results. Moreover, *Tbx21* motif enrichment and elevated T-bet protein expression in splenic T cells from anti-PD-1-treated mice (Fig. 5E,J) are consistent with *Tbx21*’s role in driving and maintaining Tex^int^ (*45*), suggesting that PD-1 blockade during priming promotes an intermediate-exhausted-like state upon recall.

As T_int_ represent a heterogenous subset, we further characterized the population using CXCR3 and CX3CR1 expression. Anti-PD-1 treatment during priming decreased T_pex_ (CXCR3^+^CX3CR1^−^), did not alter T_int_ (CXCR3^+^CX3CR1^+^), but significantly expanded KLR-exhausted (T_KLR_; CXCR3^−^CX3CR1^+^) cells upon metastatic recall (Fig. 5K). Supporting this, splenic T cells from anti-PD-1-treated mice exhibited increased KLRG1 expression and enhanced accessibility of T_KLR_-associated loci (*Klrg1*, *Fcgr2b*, *Gzmk*, *S1pr5*, *Klf2*) (Fig. 5L, Table S1). Given that CX3CR1^+^ and KLRG1^+^ T cells exhibit heightened effector capacity (*46, 47*), these findings suggest that anti-PD-1 therapy primes long-lived T cells with improved cytotoxic potential against metastatic tumors.

Together, these data suggest that PD-1 blockade during priming promotes a T_int_-like epigenetic state with T_KLR_ features during metastatic recall. As both T_int_ and T_KLR_ subsets primarily circulate rather than reside in tissues, anti-PD-1 therapy may favor a circulatory-exhausted T cell state that integrates memory-associated persistence with effector readiness, improving responsiveness to secondary encounters with metastatic tumors.

## DISCUSSION

While exhausted T cells are essential for tumor control following ICB under chronic antigen exposure, it was unknown which T cell state optimally mediates protection upon renewed antigen encounter after a period of low antigen load, such as during metastatic relapse following surgical resection. To address this, we modulated CD8^+^ T cell priming signals using anti-PD-1, IFN-β, or agonistic anti-CD40 and assessed their effects on T cell differentiation and protective immunity in a newly developed model of surgical tumor resection followed by metastatic challenge. Consistent with prior studies, IFN-β and agonistic anti-CD40 induced effector- and memory-precursor-like states, respectively. However, both interventions reduced overall survival after metastatic rechallenge, indicating that effector and memory T cell states provide suboptimal protection against metastatic lesions. In contrast, anti-PD-1 did not markedly alter early T cell differentiation but significantly enhanced protection against metastatic tumors. These results demonstrate that altering priming signals can influence durable anti-tumor immunity and suggest that intermediate-exhausted T cells are uniquely suited to provide long-term protection against metastatic disease.

Given the documented success of anti-PD-1 in both preclinical and clinical settings, its ability to induce durable, protective T cell responses was not unexpected. However, it was striking that only two doses administered during T cell priming conferred protection persisting more than 100 days after the final dose, underscoring the therapeutic potential of targeting T cell priming. Indeed, a prior study showed that anti-PD-1 therapy was most effective when given concurrently with cancer vaccination, whereas administration before or even three days after vaccination markedly diminished efficacy (*48*).

These findings point to priming as a critical determinant of long-term T cell fitness, prompting us to define the transcriptional and epigenetic states shaped by early anti-PD-1 treatment. At peak priming, transcriptional and epigenetic profiles of anti-PD-1-treated T cells closely resembled untreated controls, although scRNA-seq revealed signs of accelerated differentiation, consistent with prior work (*1, 15–17*). Supporting this, anti-PD-1-treated T cells occupied a more differentiated state in the spleen during metastatic recall. Interestingly, anti-PD-1-treated T cells at the metastatic tumor site maintained an epigenetic profile more similar to splenic T cells than untreated T cells from the tumor site. These findings suggest that anti-PD-1 accelerates initial differentiation yet stabilizes T cells in a T_int_-like state, characterized by preserved cytotoxic ability and responsiveness to ICB (*6, 7, 15*). Consistent with the maintenance of a T_int_-like phenotype, T cells from anti-PD-1-treated mice had reduced PD-1 expression within lungs at recall. Correspondingly, TBET, known to repress PD-1 expression (*34*), displayed high motif accessibility and protein expression in splenic T cells during recall, supporting its role in sustaining a T_int_-like state (*45*). Notably, a recent study in patients with melanoma correlated high TBET expression in tumor-reactive T cells with favorable responses to neoadjuvant anti-PD-1 therapy (*49*). We also identified T_KLR_ features in splenic T cells from anti-PD-1-treated mice, including increased accessibility of T_KLR_-associated loci and elevated KLRG1 and CX3CR1 expression. T_KLR_ have been described as circulatory cells with superior cytotoxic ability relative to T_term_ (*46, 47*). The induction of circulatory T_int_ and T_KLR_-like populations by PD-1 blockade may enhance clearance of transiting tumor cells and prevent metastatic seeding.

This surgical model provides a tractable platform to evaluate recall responses to metastatic lesions and dissect priming-dependent mechanisms of durable immunity. However, its translational relevance is limited by the challenge of modulating early T cell priming in patients, as cancer is diagnosed long after initial tumor-immune interactions. Nevertheless, new waves of T cell priming may be induced therapeutically through neoantigen vaccines, which are often administered in combination with anti-PD-1/PD-L1 therapy. Such combinations may synergize in a postoperative setting by inducing productive, long-lived T cell responses. Indeed, a recent study in patients with pancreatic cancer receiving adjuvant mRNA neoantigen vaccination together with anti-

PD-L1 and chemotherapy demonstrated surprisingly long-lived T cell responses (*50*). This study identified a de novo primed KLF2^+^ T cell subset, which shares features with the here identified T_int_/T_KLR_-like cell state, that correlated with long-term response. Ongoing clinical trials combining therapeutic mRNA vaccination with PD-1/PD-L1 blockade (KEYNOTE-942, KEYNOTE-603) show encouraging preliminary recurrence-free survival (*51, 52*). Together, our observations underscore the need for clinical studies testing whether blockade of the PD-1/PD-L1 pathway combined with therapies that induce T cell priming can expand durable T cell states optimally positioned to prevent recurrent disease.

## MATERIALS AND METHODS

### Cell culture and cell line generation

Melanoma was induced via topical tamoxifen application on a BRaf^CA^,Pten^−/-^,Tyr::CreER^T2^ female mouse. The tumor was harvested, minced, and incubated for 1 hour in digestion buffer containing 3mg/mL collagenase D, 1.5 mg/mL porcine pancreatic trypsin in DMEM (Gibco) for 1 hour. Cells were filtered through 100uM filter and plated. Five days later cells were transduced with lentivirus encoding LargeT antigen and allowed to grow for 5 passages. Cell line was transduced with lentivirus containing pUltra-Chili-Luc (Addgene #48688). Cells were sorted via FACS on dTomato expression to generate the Mel1 cell line. Mel1 was genotyped to confirm BRaf^V600E^ and Pten^−/-^. SNP analysis was performed by Jackson labs and confirmed 98.31% C57BL/6J and 1.69% 129S1/SvImJ. To generate Mel1.SIY, the parental cell line was transduced with a lentiviral vector pLV-Efla-IRES-Blast-SIYx3-GFP, a gift from Dr. Tom Gajewski, and selected in blasticidin (Gibco). GFP and dTomato expression were confirmed by flow cytometry. All cell lines were cultured at 37C in 5% CO_2_. Cell culture media consists of DMEM supplemented with 10% FBS (R&D Systems), 1% penicillin/streptomycin (Gibco), 10mM HEPES (Gibco), and 1X Non-essential amino acids (Gibco).

### Mice

Female C57BL/6 mice and B6 CD45.1 (Strain 002014) were purchased from Jackson labs. 2C TCR Tg CD45.1^+^ Rag2^−/-^ mice (2C mice), Rag2^−/-^ mice, and TCR-βδ^−/-^ mice were bred in house. All mice were used between 7-16 weeks of age. All animal studies were approved of by the Massachusetts Institute of Technology Committee on Animal Care.

### Tumor inoculation

Cells were harvested by trypsinization (Gibco), washed twice with PBS (Gibco), filtered through a 100μm cell strainer, and resuspended in PBS at 1×10^6^ cells/100uL for injection. Cells were injected into the flanks of mice to form subcutaneous tumors or into the tail vein for a metastatic tumor challenge.

### Tumor outgrowth

Subcutaneous tumor area measurements were performed by measuring the length and width of tumors with calipers three times per week. Tumor volume was estimated using an approximated formula for the volume of an ellipsoid: 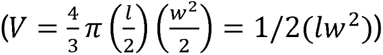.

### In vivo treatments

Anti-PD-1, clone 332.8H3, is a mouse IgG1,k antibody, which was kindly gifted by Dr. Gordon Freeman (*53–56*). Anti-PD-1 was dosed intraperitoneally (IP) at 200μg every 3 days. 100μg agonistic anti-CD40 antibody (BioXCell) was administered IP every 3 days. Recombinant IFN-β was generated in house, using a previously described protocol (*23*). The His-tagged IFN-β plasmid was gifted by Dr. Dane Wittrup. Briefly, HEK293-F cells were transfected with plasmid using PEI (Sigma-Aldrich) and allowed to grow for 6 days. IFN-β was purified from cell supernatants using a TALON Metal Affinity Resin (Takara Bio) and then buffer exchanged to PBS using Amicon filters (EMD Millipore). IFN-β was filtered and confirmed to have minimal endotoxin (<0.1 EU per dose) using the LAL Chromogenic Endotoxin Quantitation Kit (Pierce) and molecular weight was confirmed by protein gel. 0.5nM IFN-β was administered IP daily.

### Tissue harvest and dissociation

Tumor bearing mice were euthanized via CO2 asphyxiation. Spleens and lymph nodes were harvested and dissociated by mashing through 70μm cell strainer to generate a single cell suspension. Spleen suspensions were lysed with ACK lysing buffer (Gibco) for 3 minutes to remove red blood cells. Subcutaneous tumors were dissected, minced with scissors in digestion buffer (250 μg/ml Liberase (Roche) and 50 units/mL DNase (Sigma-Aldrich) in RPMI (GIBCO)) and incubated for 30min at 37C with agitation. The tumor mixture was filtered through a 70μm cell strainer to create a single cell suspension in 10mL of RPMI. To enrich for lymphocytes, 5mL of Ficoll (Sigma-Aldrich) was layered underneath the RPMI-tumor mixture. Cells were spun at 450 x g for 25 minutes with low breaks and acceleration. The interface of RPMI and Ficoll was collected into a new tube and washed prior to flow cytometry staining.

For analyses of lung tissue, mice were injected with fluorescently labeled anti-CD45 antibody retro-orbitally (CD45-IV) three minutes prior to euthanasia to differentiate cells residing in the lung vasculature and in the lung parenchyma. Lungs were dissected and dissociated using gentleMACS Octo Dissociator (Miltenyi) in a digestion buffer containing HBSS supplemented with 1mg/mL collagenaseD (Sigma-Aldrich), 50units/mL DNase (Sigma-Aldrich), 1% HEPES, and 1% FBS. The TumorImp01.01 program was used to mince tissues, which were then incubated at 37C for 30min with agitation. Dissociated lungs were passed through a 70μm filter to create a single cell suspension. ACK lysis was performed for 3 minutes, and then samples were washed with PBS prior to flow cytometry staining. For all cell sorting experiments, after a single cell suspension was obtained, CD8^+^ enrichment was performed using the CD8a+ T Cell Isolation Kit, mouse (Miltenyi Biotec). CD8^+^ cells were then stained for FACS as described above.

### Flow cytometry and FACS staining

Cells were washed in FACS buffer (PBS (Gibco), 1% FBS (R&D Systems), 2mM EDTA (Invitrogen)) prior to staining. Cells were stained for 15 minutes at 4C with Fixable Viability Dye eFluor 780 or eFluor 506 (eBioscience) and αCD16/CD32 (clone 93, BioLegend) to identify live cells and to prevent non-specific antibody binding. Cells were washed with FACS buffer and stained at 4C for 30 minutes for surface markers using fluorescently labeled antibodies diluted in FACS buffer at specified dilutions (table S2). Cells were washed twice in FACS buffer and directly analyzed or were fixed for intracellular staining and/or for analysis the following day. Fixation was performed using the Foxp3 Transcription Factor Fixation/Permeabilization buffer (eBioscience) following the manufacturer’s instructions. For intracellular staining, fluorescently labeled antibodies were suspended in 1X Permeabilization Buffer (eBioscience) and stained for a minimum of 30 minutes at 4C. To obtain cell counts, Precision Count Beads (BioLegend) were added to samples following manufacturer’s instructions after all staining was complete. Flow cytometry was performed on a LSR Fortessa cytometer (BD) or a FACSymphony cytometer (BD). For cell sorting, a FACSAria (BD) was used.

SIY-reactive CD8^+^ T cells were isolated via FACS and collected in T cell media (RPMI (Gibco), 10% FBS (R&D Systems), 1% penicillin-streptomycin (Gibco), 10mM HEPES (Gibco), 1X Non-essential amino acids (Gibco), 1mM Sodium Pyruvate (Gibco), 2-mercaptoethanol (Gibco)). Staining for SIY-reactive T cells was performed using tetramer reagents generated from biotinylated monomers (NIH Tetramer Core Facility) and streptavidin-R-phycoerythrin conjugate (Invitrogen), following the NIH Tetramer Core guidelines. All flow cytometry analysis was performed using FlowJo v10.5.3 software (TreeStar).

### Single cell RNA sequencing and single cell TCR sequencing using Seq Well

TdLNs were harvested, single cell suspensions were created, enriched for CD8^+^ cells, and stained for FACS as described above. SIY-tetramer^+^ CD8^+^ T cells were sorted via FACS and collected in T cell media, as described above. Single-cell RNA sequencing was performed on the Seq-Well platform with second-strand chemistry as previously described (*35, 57*). Libraries were dual-barcoded and amplified using Nextera XT (Illumina) and sequenced on an Illumina Novaseq 6000. Paired TCR and sequencing alignment was performed as previously described (*58*). In summary, biotinylated *Tcrb* and *Tcra* probes in conjunction with magnetic streptavidin beads were used to enrich for TCR transcripts from the whole transcriptome amplification product from each library and then were further amplified using V-region primers and Nextera sequencing handles. TCR libraries were sequenced on an Illumina MiSeq.

### scRNA-seq data analysis

Raw sequencing data was demultiplexed and processed using the Drop-seq pipeline (*59*). Reads were aligned to the mm10 genome and collapsed by cell barcode and unique molecular identifier (UMI). We filtered out genes that were detected in less than 10 cells and cells in which fewer than 1,000 unique features were detected. Analysis was performed in Seurat (*60*) using standard functions. Briefly, data were normalized using SCTransform and linear dimensionality reduction was performed using RunPCA. The number of principal components to use was determined by elbow plot. Clustering was performed using FindNeighbors and FindClusters. Cell typing was performed based on top genes from FindAllMarkers. Spearman correlation was calculated based on pseudobulk values using AggregateExpression. Differential gene expression was performed using MAST (*61*). Gene set expression analysis was performed using fgsea package (*62*) and the Hallmark gene sets.

### scTCR-seq analysis

Cells with beta chain sequences were retained for analysis (i.e., alpha chains were dropped). Cells with ambiguous beta chains, defined by having >1 beta chain with >1 UMI per beta chain, were also dropped. Beta chains with only 1 UMI were removed. The C gene was set to a constant value (TRBC) for all chains. Clone calling was performed using combineTCR from scRepertoire (*63*) and the CTstrict clone definition was used for all downstream analysis (identical V, D, J, and C genes and JUNCTION at the nucleotide level). Clonal diversity indices were computed using Alakazam (*64*). The Morisita index was computed using the clonalOverlap function in scRepertoire.

### Tumor resection surgery

Tumor resection surgery was performed on mice bearing Mel1.SIY tumors. Mice were given 1mg/kg buprenorphine extended-release subcutaneously as an analgesic. Hair was removed around tumor region. Mice were anesthetized with inhaled isofluorane. Surgical area was sterilized and the tumor with attached skin flap were removed. Incisions were closed using wound clips. Mice were monitored daily for 3 days following surgery to confirm appropriate recovery and wound healing. Wound clips were removed ten days after surgery when incision had fully closed.

### Blood collection and analysis

Blood was collected from mice via submandibular vein puncture in a tube containing 50uL heparin (Sigma-Aldrich). Red blood cells were removed with ACK lysing buffer (Gibco). Cells were washed with FACS buffer and stained as previously described.

### 2C T cell adoptive transfer

2C mice were euthanized and lymph nodes and spleens were harvested and turned into single-cell suspensions as described above. Cells were washed and resuspended in PBS (Gibco). 1 million cells were transferred retro-orbitally into day 3 Mel1.SIY flank tumor-bearing mice. Mice were treated with therapy as described. Four days after transfer, primary recipients were euthanized and TdLNs harvested. Congenically-marked T cells were analyzed via flow cytometry or were isolated via FACS for secondary transfers. For secondary transfers, live CD45.1^+^ T cells were sorted, and 20,000 cells were transferred retro-orbitally into secondary recipients, which were either day 4 Mel1.SIY tumor-bearing Rag2^−/-^ mice or naïve TCR-βδ^−/-^ mice.

### In vivo cytotoxicity assay

This assay was adapted from a previously published protocol (*65*). Briefly, spleens collected from naïve CD45.1^+^ mice and dissociated to single cells as described above. Cells were stained with either CellTrace Violet (Life Technologies) or CellTrace CFSE (Life Technologies) following manufacturer’s instruction. Cells were then loaded with 10μg/mL SIY peptide for 2 hours in T cell media. Unloaded cells were used as control. Cells were washed and counted and then mixed at a 50:50 ratio of SIY-loaded to unloaded. 5×10^6^ cells were transferred retro-orbitally into day 6 Mel1.SIY tumor-bearing mice. 24 hours later, spleens of recipient mice were harvested and prepared for flow cytometry. The ratio of peptide-pulsed and unpulsed CD45.1^+^ cells was determined and normalized to the ratio found in naïve control mice. Percent lysis was calculated 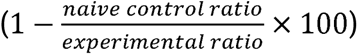.

### Single cell ATAC sequencing

Mice were inoculated with Mel1.SIY flank tumors and left untreated or treated with anti-PD-1 therapy at day 0 and day 3 after tumor inoculation. On day 6, mice were euthanized and TdLNs harvested. TdLNs were physically dissociated by mashing in a scored plate and incubated with a digestion buffer (HBSS supplemented with 1mg/mL collagenaseD (Sigma-Aldrich), 50units/mL DNase (Sigma-Aldrich), 1% HEPES, and 1% FBS) for 15min at 37C before being passed through a 70μM cell strainer. CD8^+^ enrichment was performed and SIY-reactive T cells were sorted as previously described. Alternatively on day 6, tumors were surgically removed and mice allowed to recover for 7-8 weeks. Mice were then rechallenged with Mel1.SIY injected IV. 14 days later (on day 59 after initial flank tumor inoculation), mice were euthanized 3 minutes after retro-orbital injection of anti-CD45-IV. Spleens and lungs were prepared for FACS as described above. CD45-IV negative, SIY-tetramer^+^ T cells were sorted from spleen and lung. The 10X Genomics Nuclei Isolation Protocol (CG000169 RevE) for low cell input was followed for nuclei isolation. In brief, cells were washed and then incubated in lysis buffer (10mM Tris-HCl, 10mM NaCl, 3mM MgCl2, 0.1% Tween-20, 0.1% IGEPAL CA-630, 0.01% Digitonin, 1% BSA) for 3 minutes. Cell lysis was confirmed with trypan blue staining. Nuclei were submitted to the MIT BioMicro Center for library generation and sequencing on the Element AVITI24.

### Single cell ATAC analysis

Raw sequencing reads were processed with cellranger-atac (v2.1.0). The reference mouse genome used was GRCm39/mm39 (https://www.ncbi.nlm.nih.gov/datasets/genome/GCF_000001635.27/). Further analyses were performed in R (v4.4.1) using the ArchR package (v1.0.3). A blacklist region file for mm39 was obtained from https://dev.bedbase.org/bedset/excluderanges. The mm39 genome annotation was retrieved from R package ‘BSgenome.Mmusculus.UCSC.mm39’ and gene annotations were retrieved from R package ‘TxDb.Mmusculus.UCSC.mm39.refGene’. Initial arrow files were created with the createArrowFiles command using options minTSS=1 and minFrags=1000. Low quality cells were filtered after inspection of the quality control images of the number of nuclear fragments versus TSS enrichment using the tss.plot function. In particular, we used the log10 nuclear fragment cutoffs 3.1, 3.5, 3.5, 3.5, 3.5, 3.4 and the TSS enrichment cutoffs 8, 8, 12, 12, 18, 18 for the corresponding samples D6-TdLN-U, D6-TdLN-P, D59-L-U, D59-L-P, D59-S-U, D59-S-P. Fragment size distribution images were created with the plotFragmentSizes function. From the filtered cells we removed doublets by using the addDoubletScores function and the filterDoublets function with default parameters. The raw fragment counts were then transformed and normalized by using the Latent Semantic Indexing method. The obtained data matrix was subjected to singular value decomposition and the singular vectors were subsequently used for further downstream analyses. In particular, singular vectors were used for making a KNN graph that was then used for Louvaine clustering to identify groups of cells from all six samples with similar accessible chromatin regions. To visualize the transformed fragments a 2-D UMAP embedding was calculated with the singular vectors as input.

Differential accessible regions (DAR) between pre and post anti-PD-1 treatment of the same tissue were determined by first using the addReproduciblePeakSet function and then the getMarkerFeatures function from the ArchR package. We further determined the closest gene to each DAR by using the closest function from bedtools (*66*).

To determine if certain TF motifs are enriched in the DARs we analyzed all six samples together. First, we redid the DAR calling on all samples. Second, we determined TF motifs in DARs by using the addMotifAnnotations routine using the available ‘cisbp’ (*67*) dataset. Last, we called the peakAnnoEnrichment function and set a FDR cutoff of 0.1 and demanded a log2 foldchange of at least 1 corresponding to an enrichment of at least twofold.

### Projection of bulk ATACseq data from Beltra et al. 2020 (*45*)

Raw data was obtained from GEO with accession number GSE149879. The FastQ files were then processed with the nfcore atacseq pipeline to get aligned reads to the mm39 genome. The resulting bam files were then used to create a count matrix that indicates how many ATACseq reads per sample are mapped to any of our identified scATACseq peaks from our data. The count matrix was used as input for the projectBulkATAC function that projected the Bulk data to our 2D UMAP embedding.

### Statistical analysis

All statistical analyses were performed in GraphPad Prism v10. All data are shown as means ± SEM, unless otherwise indicated. Statistical significance of flow cytometry experiments was determined by Mann-Whitney test comparing two groups and by one-way ANOVA or Kruskal-Wallis test comparing multiple groups. For experiments with low sample sizes (n<10 per group), Shapiro-Wilk test was used to determine normality. If passed, experiments with more than two groups were analyzed by one-way ANOVA with a posttest of Holm-Sidak’s multiple comparisons test was used and experiments with only two groups were analyzed by unpaired T test. If the dataset was determined to not follow a normal distribution and had more than two experimental groups, a Kruskal-Wallis test was performed with a posttest of Dunn’s multiple comparisons test, comparing treated samples to the untreated controls. If an experiment had only two experimental groups and did not follow a normal distribution, a Mann Whitney test was performed. For tumor outgrowth studies, 2-way ANOVA followed by Dunnett’s multiple comparisons test was performed. Analyses of survival studies was performed with a log-rank (Mantle-Cox) test, comparing specific groups of interest. For each experiment, two to four independent repeats were performed and combined; *n* numbers are indicated in figure captions. *p<0.05, **p<0.01, ***p<0.001, ****p<0.0001.

## Acknowledgments

We would like to thank Dr. Gordon Freeman for generously providing the anti-PD-1 used in these studies; Melissa Duquette for mouse colony maintenance and laboratory support; Paul Thompson and Judy Teixeira for administrative support; and the Koch Institute Flow Cytometry core and the MIT BioMicro Center for technical support.

## Author contributions

Conceptualization: TD, SS

Methodology: TD, SS

Investigation: TD, SM, ZJR, VB, MYC, YJZ, DM, LP, YJ, AR

Visualization: TD, SM, ZJR

Resources: TD, FC, EL

Funding acquisition: SS

Supervision: SS, EL, KDW, JCL, AM,

Writing – original draft: TD

Writing – review & editing: TD, ZJM, FC, VB, SS

