## Supplementary figures and images for "PD-1 blockade during T cell priming enhances long-term protection against metastatic tumors by epigenetically tuning T cell exhaustion"

### Fig S1

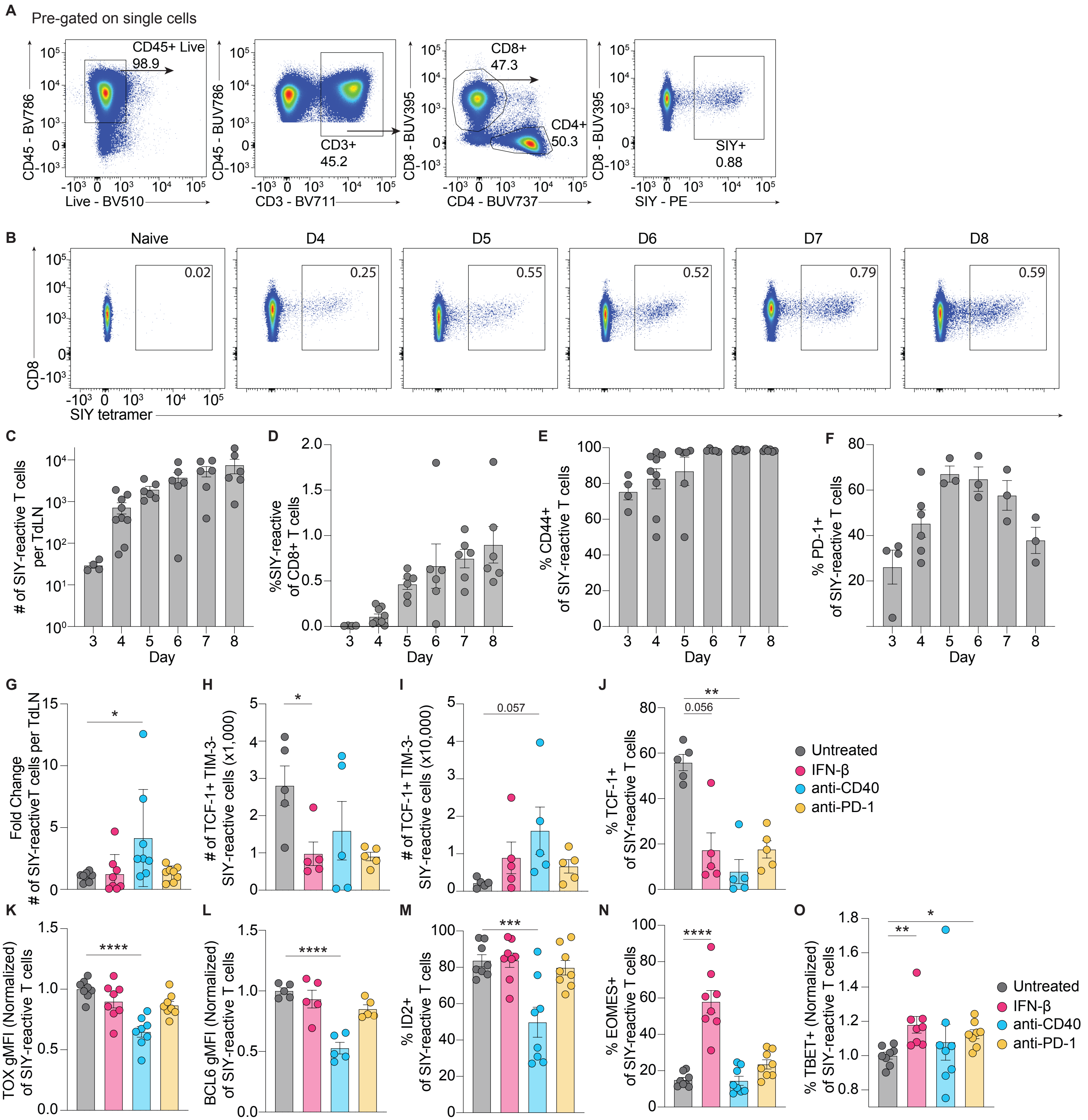

### Fig S2

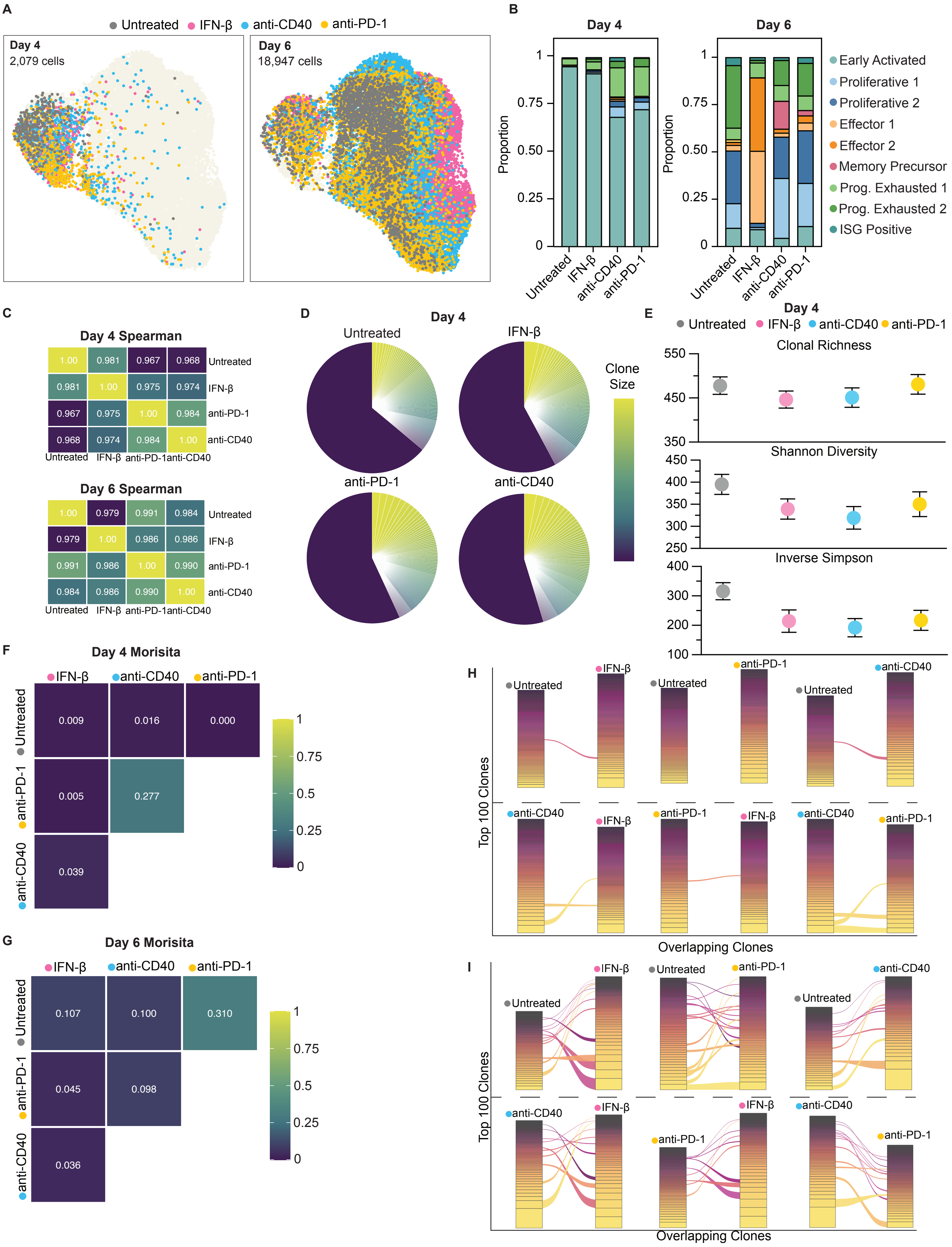

### Fig S3

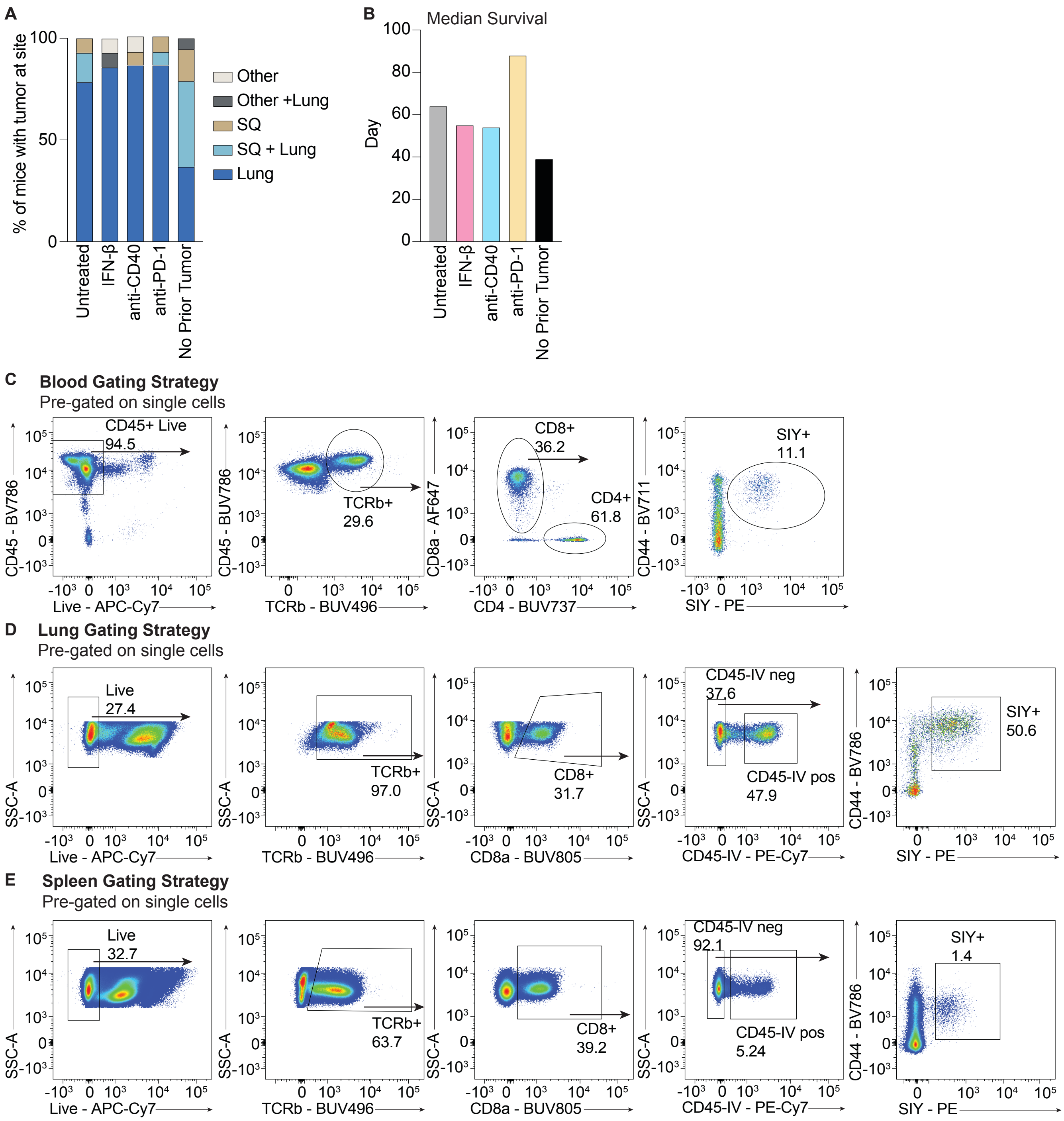

### Fig S4

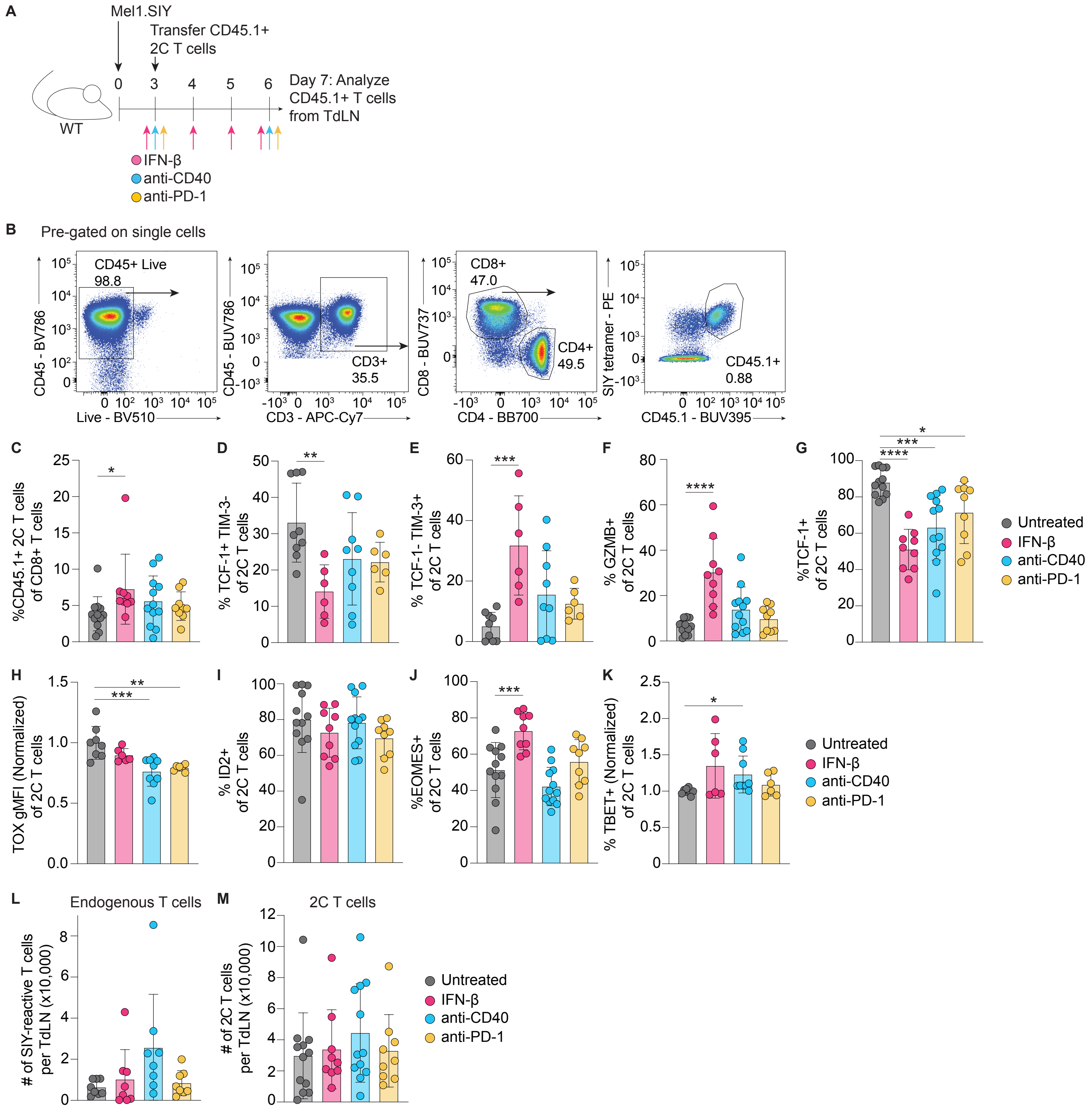

### Fig S5

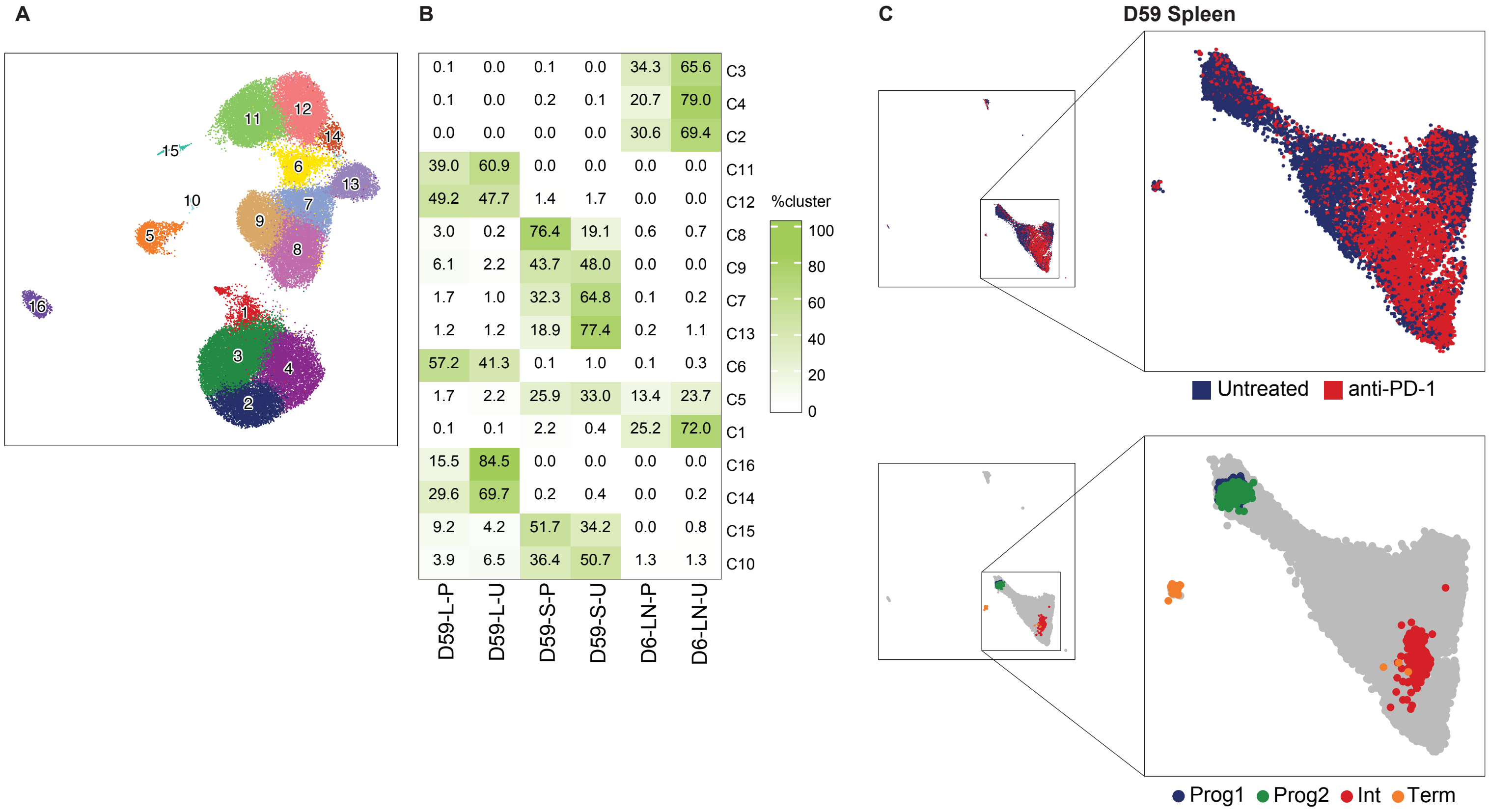
